# A validated strategy to infer protein biomarkers from RNA-Seq by combining multiple mRNA splice variants and time-delay

**DOI:** 10.1101/599373

**Authors:** Rasmus Magnusson, Olof Rundquist, Min Jung Kim, Sandra Hellberg, Chan Hyun Na, Mikael Benson, David Gomez-Cabrero, Ingrid Kockum, Jesper Tegnér, Fredrik Piehl, Maja Jagodic, Johan Mellergård, Claudio Altafini, Jan Ernerudh, Maria C. Jenmalm, Colm E. Nestor, Min-Sik Kim, Mika Gustafsson

**Affiliations:** Bioinformatics, Department of Physics, Chemistry and Biology, Linköping University, Linköping, Sweden; Department of Applied Chemistry, College of Applied Sciences, Kyung Hee University, Yong-in 446-701, Republic of Korea; Department of Clinical and Experimental Medicine, Linköping University, Linköping, Sweden; Department of Neurology, Institute for Cell Engineering, Johns Hopkins University School of Medicine, Baltimore, MD, USA; Centre for Personalised Medicine, Linköping University, Linköping, Sweden; Navarrabiomed, Complejo Hospitalario de Navarra, Universidad Pública de Navarra, IdiSNA, 31008 Pamplona, Spain; Department of Clinical Neuroscience, Center for Molecular Medicine, Karolinska Institute, 171 77, Stockholm, Sweden; Biological and Environmental Sciences and Engineering Division, Computer, Electrical and Mathematical Sciences and Engineering Division, King Abdullah University of Science and Technology (KAUST), Thuwal 23955–6900, Saudi Arabia; Unit of Computational Medicine, Department of Medicine, Solna, Center for Molecular Medicine, Karolinska Institutet, Stockholm, Sweden; Science for Life Laboratory, Solna, Sweden; Department of Neurology, Linköping University, Linköping, Sweden; Department of Automatic Control, Linköping University, Linköping, Sweden; Department of Clinical Immunology and Transfusion Medicine and Department of Clinical and Experimental Medicine, Linköping University, Linköping, Sweden; Department of New Biology, Daegu Gyeongbuk Institute of Science and Technology, Daegu 711-873, Republic of Korea

## Abstract

**Background:** Profiling of mRNA expression is an important method to identify biomarkers but complicated by limited correlations between mRNA expression and protein abundance. We hypothesised that these correlations could be improved by mathematical models based on measuring splice variants and time delay in protein translation.

**Methods:** We characterised time-series of primary human naïve CD4^+^ T cells during early T-helper type 1 differentiation with RNA-sequencing and mass-spectrometry proteomics. We then performed computational time-series analysis in this system and in two other key human and murine immune cell types. Linear mathematical mixed time-delayed splice variant models were used to predict protein abundances, and the models were validated using out-of-sample predictions. Lastly, we re-analysed RNA-Seq datasets to evaluate biomarker discovery in five T-cell associated diseases, validating the findings for multiple sclerosis (MS) and asthma.

**Results:** The new models demonstrated median correlations of mRNA-to-protein abundance of 0.79-0.94, significantly out-performing models not including the usage of multiple splice variants and time-delays, as shown in cross-validation tests. Our mathematical models provided more differentially expressed proteins between patients and controls in all five diseases. Moreover, analysis of these proteins in asthma and MS supported their relevance. One marker, sCD27, was clinically validated in MS using two independent cohorts, for treatment response and prognosis.

**Conclusion:** Our splice variant and time-delay models substantially improved the prediction of protein abundance from mRNA data in three immune cell-types. The models provided valuable biomarker candidates, which were validated in clinical studies of MS and asthma. We propose that our strategy is generally applicable for biomarker discovery.

## Introduction

A key problem in genome medicine is to find reliable disease biomarkers and therapeutical targets. An important reason is that common diseases involve thousands of proteins across multiple cell types. Proteins are regarded as optimal biomarkers as they are the main drivers of the crucial functions necessary for life, and thus directly connected to patho-physiological processes[1]. Furthermore, many proteins can be readily measured in biological fluids. However, proteome-wide analyses are difficult to perform in clinical studies due to the large quantities of material needed. On the other hand, gene expression profiling can be performed using a range of techniques, such as microarrays or RNA-sequencing. Another advantage of using mRNA expression as a core vehicle for biomarker discovery is that mRNA profiling can be performed even if only samples of limited amount, like biopsies, are available.

Combinations of mRNAs can have high diagnostic efficacy in multiple diseases[2, 3]. An ideal solution could therefore be to perform mRNA profiling to identify protein biomarkers that are needed for diagnosing and subtyping of diseases, as well for the personalisation and monitoring of treatments. However, this approach is complicated by the low correlation between mRNA and protein expression[4-7], which can be tackled with different strategies[8, 9]. The discrepancy between mRNA and protein abundance is due to several factors, including but not limited to differences in the rates of translation and degradation between proteins and cell-types[10]. Moreover, the data resolution of mRNA splice variants and protein isoforms further complicates such analyses, as in the cases of unequal contribution of individual splice variants to the production of a given protein[11], and cell-type specific differences in splice variant use[12].

Thus, the inability to predict protein abundance from mRNA abundance represents a major limitation in biomarker discovery. To this end, we developed a novel method to infer protein levels from mRNA expression data. Our procedure was derived by experimentally analysing early human T helper 1 (T_H_1) differentiation and constructing a machine learning modelling approach for time-series RNA-Seq and proteomics data from a dynamical perturbation of the cell-type of interest. T_H_ differentiation is an optimal model system to dissect the relationship between mRNA and protein as (i) primary human naïve T_H_ (NT_H_) cells can be isolated in high purity and large quantity from human blood (ii), all NT_H_ cells are synchronised in the G_1_ phase of the cell cycle, further reducing inter-cell heterogeneity[13] and (iii) easy access to large quantities of material allows changes in mRNA and associated protein abundance to be assayed over time[14]. Moreover, T_H_ cells are important regulators of immunity and thereby associated with many complex diseases, and T_H_1 differentiation itself is pathogenetically relevant in several diseases[15]. The utilised models were based on a time-delayed linear model between mRNA splice-variants of the same gene and protein levels. We generalised the model by applying it onto recent data from human regulatory T (T_reg_) cell and murine B cell differentiation. By combining the strength of time-series analysis and RNA-sequencing, we were able to increase median mRNA-protein correlations significantly from the initial 0.21 to 0.86. Next, we showed the potential clinical usefulness of our derived models by detecting potential biomarkers in five complex diseases. This application revealed significantly more predicted biomarkers than by using off-the-shelf methods for RNA-Seq data analysis only. Analysis of these predicted proteins in asthma and MS supported their biological relevance. Finally, we validated one of the predicted biomarkers using two independent multiple sclerosis cohorts, which showed a remarkably better stratification between patients and controls than any of our previously reported protein biomarkers. The application of our approach to multiple different cell types, species and diseases shows its general applicability to increase the power of RNA-Seq based studies for biomarker discovery.

## Results

### A significant portion of T-cell genes showed diverse correlations between RNA splice variants and proteins

In order to generate accurate mRNA and protein models, taking into account the major factors of time-delay and splice variant usage, we first developed a model analysing early T-helper type 1 (T_H_1) differentiation. This was done by performing time-series RNA-sequencing and mass-spectrometry proteomics of primary human NT_H_ cells (**Figure 1A, S1, S2**). RNA-seq (> 40× 10^6^ reads per sample) and proteome profiling was performed to detect differentially expressed mRNA splice variants and proteins at six time points from 30 minutes to five days of T_H_1 differentiation (**Figure 1A, S1, S2**). This approach detected 6909 proteins, of which 4920 could be mapped to genes expressed in the RNA-Seq data. As expected, a significant fraction of the genes showed a significant positive correlation between mRNA and protein levels (n=407, expected 123 out of 4920 proteins, binomial test P<10^−93^) during T_H_1 cell differentiation. Interestingly, a significant fraction of negatively correlated genes was also observed (n=205, expected 123, P<10^−11^) (**Figure 1B, Table S1**). Remarkably, the overall median Pearson correlation (rho) between mRNA and protein was only 0.21. We hypothesised that this could depend on variable correlations between mRNA splice variants of each gene and the protein it encoded. Indeed, we found both positive and negative correlations between splice variants and their corresponding proteins (binomial test for enrichment of significant negative correlation P<1.3 × 10^−3^, odds ratio= 1.48). For example, the known T_H_ cell associated genes, *IL7R* and *STX12*[16] contained multiple splice variants, of which several were positively or negatively correlated to their corresponding protein levels (**Figure 1C**). Given the large variation in correlation between different splice-variants of a given gene and its corresponding protein, we proceeded to construct predictive splice-variant models of protein abundance.

**Fig 1.**
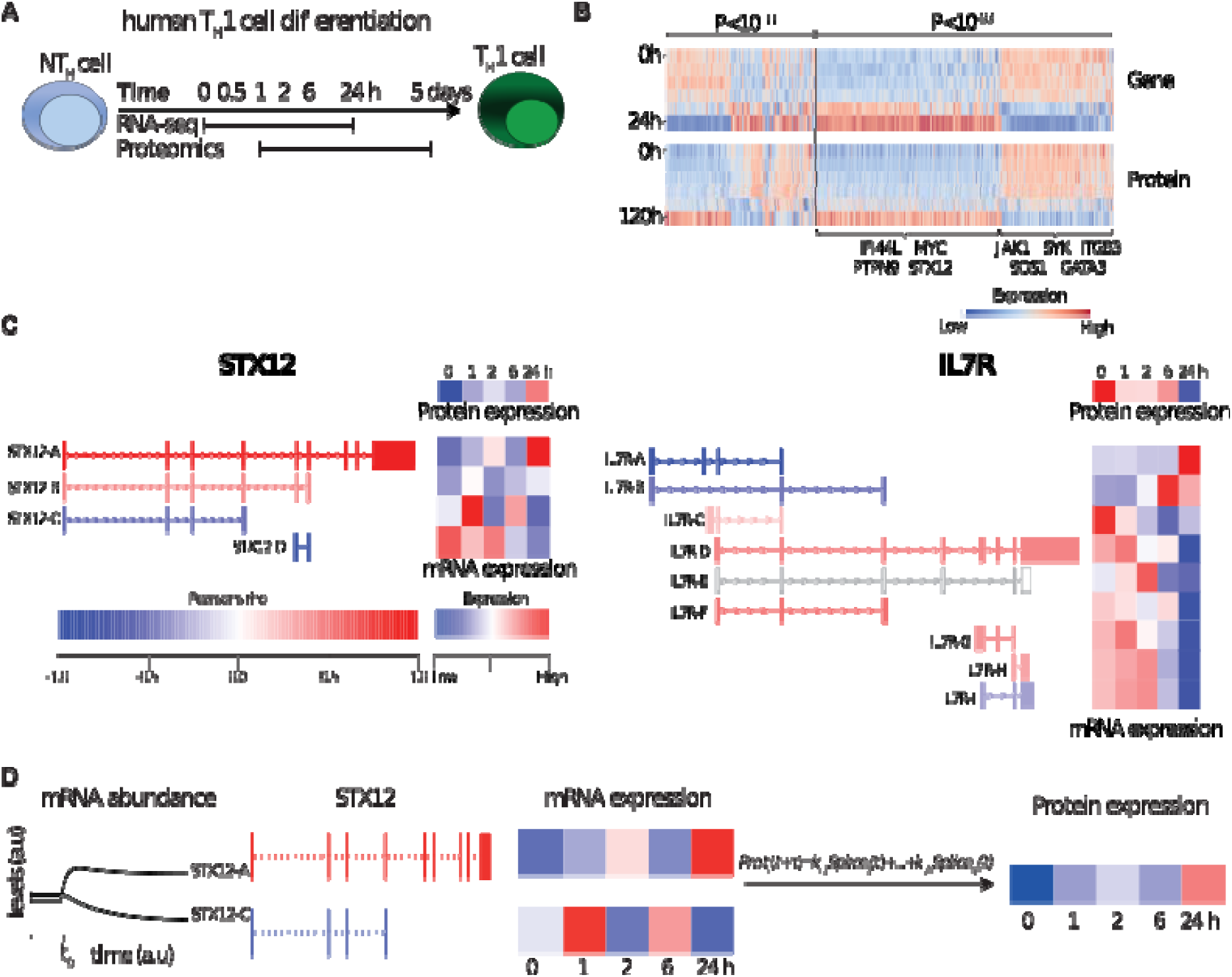
RNA-Seq and mass-spectrometry analysis of T_H_1 differentiation revealed highly variable correlations. (A) Experimental design. (B) Heat map of transcript and protein abundance dynamics in genes that show significant negative (left) and positive (right) correlations. (C) Examples of transcript splice variants showing that both *STX12* (left) and *IL7R* (right) were significantly negatively and positively correlated with protein levels. (D) Illustration of the modelling procedure for resolving the poor correlation, using STX12 as an example.

### A linear model combining the expressions of multiple splice variant transcripts showed substantially stronger correlations with protein abundance than individual transcripts

In order to construct generally applicable and predictive mRNA-to-protein models, we applied a simple linear relation between the protein abundance of a gene and its associated mRNA splice-variants. Furthermore, we allowed for different translation times for each gene. Firstly, we used a cross-validated L1 penalised linear regression model to favour simple models using single splices without any time-delays (Methods, **Figure 1D**). The rationale for the L1 penalty was to effectively remove splice variants that carry little or no predictive power over protein abundance. This simple model resulted in a median gene-protein correlation of *rho*_TH1_ = 0.86 (**Figure 2A)**, far in excess of previously reported gene-protein prediction models in mammals[5, 7, 10, 11]. Likewise, we also trained similar models for two existing mRNA-protein time-series datasets with similar results, that is from human T_REG_ cells[14](*rho*_TREG_ = 0.79) and mouse B cells (GSE75417) (*rho*_Bcell_ = 0.94) (**Figure 2A**). In order to test whether the increase in correlation was due to the incorporation of negatively correlating splice variants, multiple transcripts, or time-delay we also constructed such models without each of these effects. Importantly, our model out-performed models with one splice variant for each gene (*rho*_TH1_ = 0.71, *rho*_TREG_ = 0.44, *rho*_Bcell_ = 0.52), and models using multiple transcripts but without a time delay (*rho*_TH1_ = 0.74, *rho*_TREG_ = 0.69, *rho*_Bcell_ = 0.45) (**Figure 2B-C**), thus demonstrating that both multiple dynamical splice variants and time delay are needed for optimal performance. In order to define the optimal time-delays between splice-variants and proteins, we analysed the time delay distributions and found it to have a mean of 8h 17 min, 6h 18 min and 8h 49 min for T_H_1, T_REG_ and murine B cells, respectively. The detailed parameters of our models are fully displayed in Table S1. Next, by using cross-validation we confirmed that our models could do out-of-sample prediction significantly better than gene expression-based models of protein abundance (binomial test; P_TH1_= 10^−152^, P_TREG_= 10^−247^, P_mice B_= 10^−59^), and better than static splice-variant models which did not include time-delays (P_TH1_=10^−1459^, P_TREG_= 10^−8^, P_mice B_= 5× 10^−4^, Fig. 2B). To evaluate mRNA-protein associations in steady state across tissues, we used mRNA expression data from the human protein atlas[17]. We found only marginal improvements by using splicing information in the multi-tissue models with respect to what had previously reported in the literature[5](rho_ProtAtlas_= 0.27, see **Figure S3)**. This lack of correlation may be explained by the lack of dynamic data, and by the presence of different cell types, and we speculate that differences in splice variant specificity between tissues effectively hinders this type of models. In further support of cell type specificity, we found only marginal correlations (*rho* = 0.09) when comparing the correlation coefficients of our two T-cell datasets of T_H_1and T_REG_ cells. Thus, a common unifying model for many cell-types remains a challenge (**Table S1**). In summary, we have revealed that by using a simple linear model of mRNA splice variants and time delay, we could predict protein abundances accurately.

**Fig 2.**
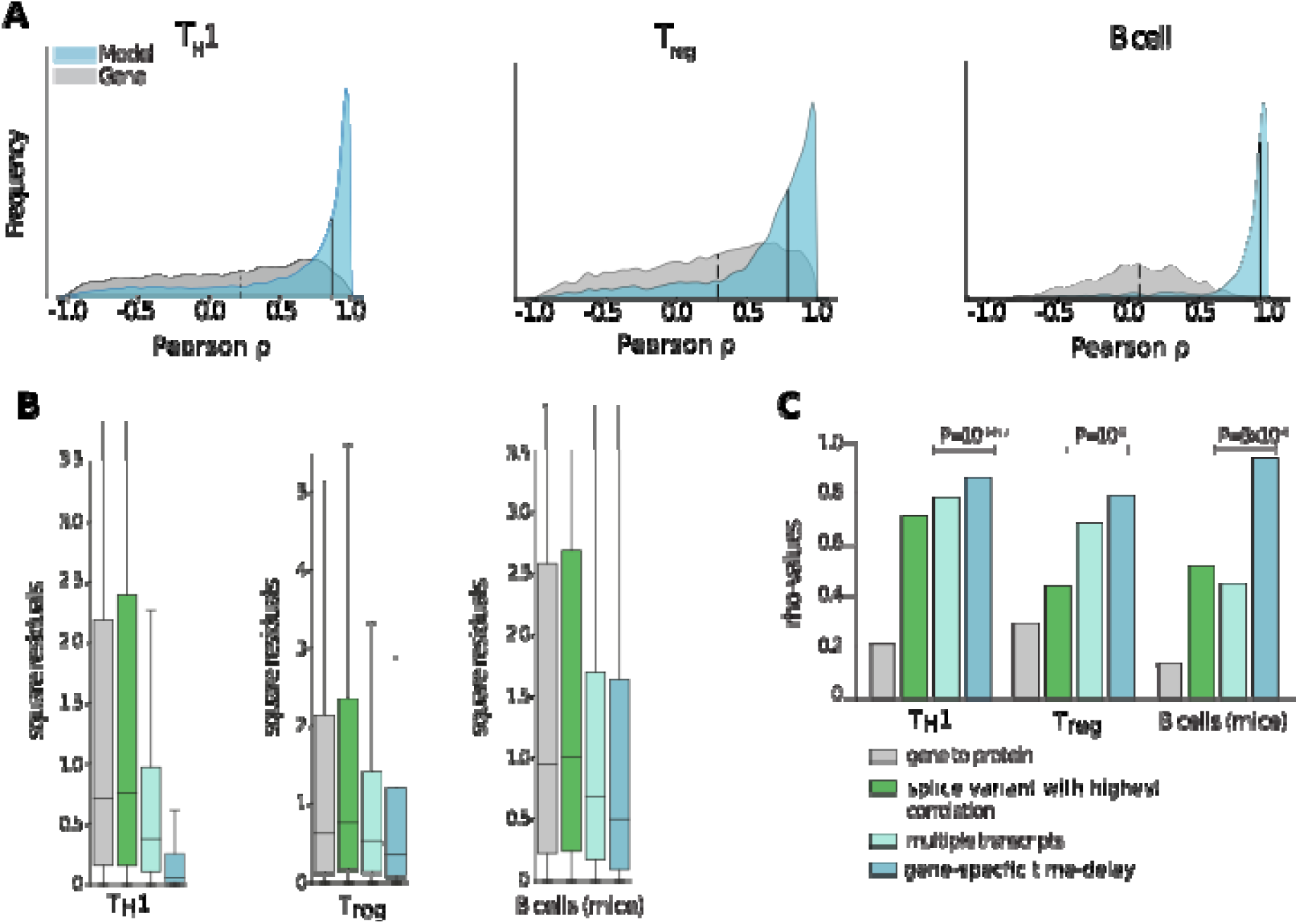
Multiple transcripts and time-delays increased mRNA and protein correlations significantly in multiple cell-types. (A) Gene/protein Pearson correlations in T_H_1 (left), T_reg_ (middle left), and murine B-cell (middle right) differentiation. In the histogram, the grey curve shows the correlation distribution when the sum of all splice variant expressions of a transcript [4] is used to quantify mRNA abundance (median: dashed line), while in the blue histogram our time-delayed multiple splice variant based models are used (medians: solid lines at 0.86, 0.79, and 0.94 for T_H_1, T_reg_ and murine B-cells, respectively). Only cross-validated protein predictions are shown for the proteins for which the null-model could be rejected. (B) Out-of-sample cross validation prediction of the three models. Aiming to quantify the predictive power of each added input to the model, we observed that a linear model with gene-specific time-delays was the model that generated predictions with the smallest sum of squared residuals. (C) Median correlation coefficients (rho) for different mathematical protein prediction models derived from RNA with increasing protein abundance correlations. P-values were derived from predictions using leave-one-out cross-validation.

### Applying the model to clinical datasets revealed potential biomarkers which were validated in multiple sclerosis and asthma

Lastly, we aimed to test the potential usefulness of our derived models for the identification of protein biomarkers by applying them on available RNA-Seq datasets from human total CD4^+^ T cells. We found data-sets for five different diseases[18-21]; asthma, allergic rhinitis, obesity-induced asthma, pro-lymphocytic leukaemia, and multiple sclerosis (MS), as well as corresponding controls. Because our models correlated well to protein abundances, we hypothesised that differential expression tests using the predicted proteins between patients and controls to be more sensitive than testing directly on the mRNA expression for all splice variants individually. Indeed, we observed that the fraction of nominally differentially expressed genes was higher than using an individual differential expression analysis for all ten comparisons (binomial P< 9.8 ×10^−4^) (**Figure 3A**). Moreover, we consistently observed a higher enrichment for the T_H_1 model compared to the T_REG_ model (P<0.03), with the highest enrichments in MS and asthma. We therefore proceeded to use our T_H_1 model on MS and asthma.

**Fig 3.**
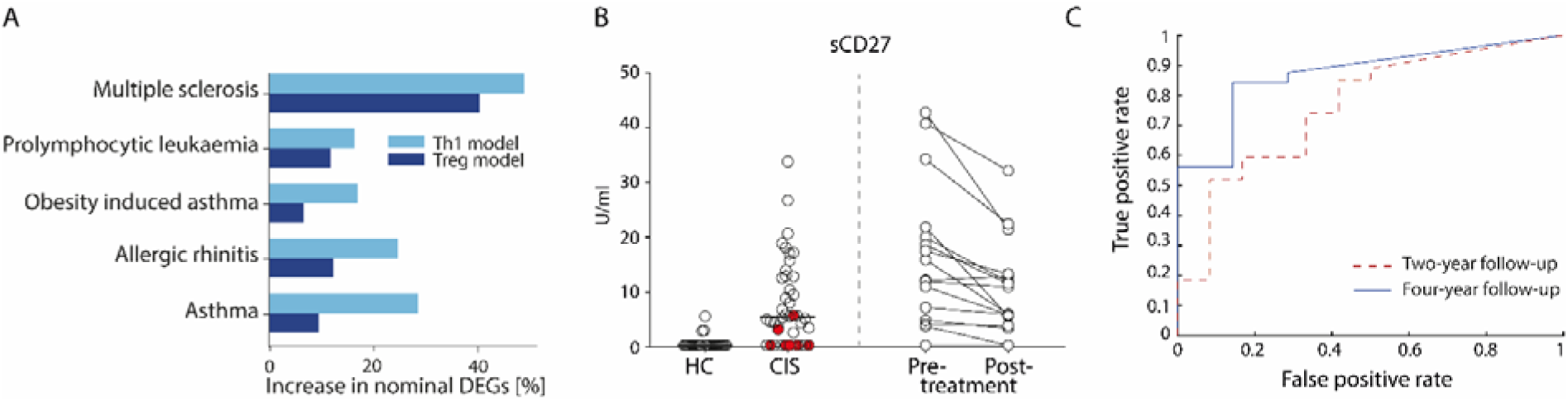
Proteins models led to the discovery of new potential biomarkers of complex diseases that were validated in multiple sclerosis (MS). (A) Differential predicted protein (PP) analysis of five diseases using the T_H_1 (light blue) and T_reg_ (dark blue) models showed higher fraction of nominally significant genes than that of normal differential gene expression tests. (B) Validation of PP from early MS (clinically isolated syndrome (CIS)) *vs* healthy controls (HC) and pre *vs* post one-year treatment with Natalizumab. Validation is performed measuring sCD40 in cerebrospinal fluid (CSF) and stratifying on phenotyping. Left plots show healthy controls vs CIS showing patients with no evidence of disease activity (NEDA) at four years treatment with filled circles. (C) Receiver operating curve using sCD27 concentration as a single prognostic marker of NEDA at four (solid line) and two years (dashed line) after CIS.

For MS, we found 20 genes with FDR<0.05, of which none could be found by testing for differential expression on the mRNA expression data directly (**Table S2**). Interestingly, eight of the 20 proteins had previously been associated with MS (**Figure 4**)[22-31]. In order to further justify the relevance of the added proteins as potential biomarkers, we proceeded to study three secreted proteins that our model predicted to be differentially expressed in the MS dataset (Annexin A1, sCD40L and sCD27). Notably, these proteins have been associated with MS previously[22, 23, 25]. We analysed if cerebrospinal fluid (CSF) levels of these proteins related to clinical outcome and immunomodulatory treatment in two independent cohorts, namely newly diagnosed MS patients (clinically isolated syndrome (CIS) and relapsing/remitting MS, n=41) *vs* healthy controls (HC, n=23), and response to Natalizumab treatment in relapsing remitting MS patients (see supplementary notes, n=16). In both cohorts, only sCD27 was present at a detectable level, while Annexin A1 and sCD40L were not. Analysis of all patients (n=57) *vs* HC (n=23) showed high separation (AUC=0.88, non-parametric P=3.0 × 10^−8^, **Figure 3B**), and treatment with Natalizumab reduced the sCD27 levels by 34% (P=4.9 × 10^−4^). Notably, sCD27 levels at baseline of newly diagnosed MS patients were able to predict disease activity after four years follow up (AUC= 0.87, P=1.2 × 10^−3^, **Figure 3B**), which was a stronger prediction than that of all our previously reported 14 biomarkers[32]. Taken together, using the splice variants-to-protein model we were able to *uniquely* identify and validate biomarkers of MS in an independent patient cohort, while these genes could not be discovered using previous state-of-the-art test for differential gene expression.

**Fig 4.**
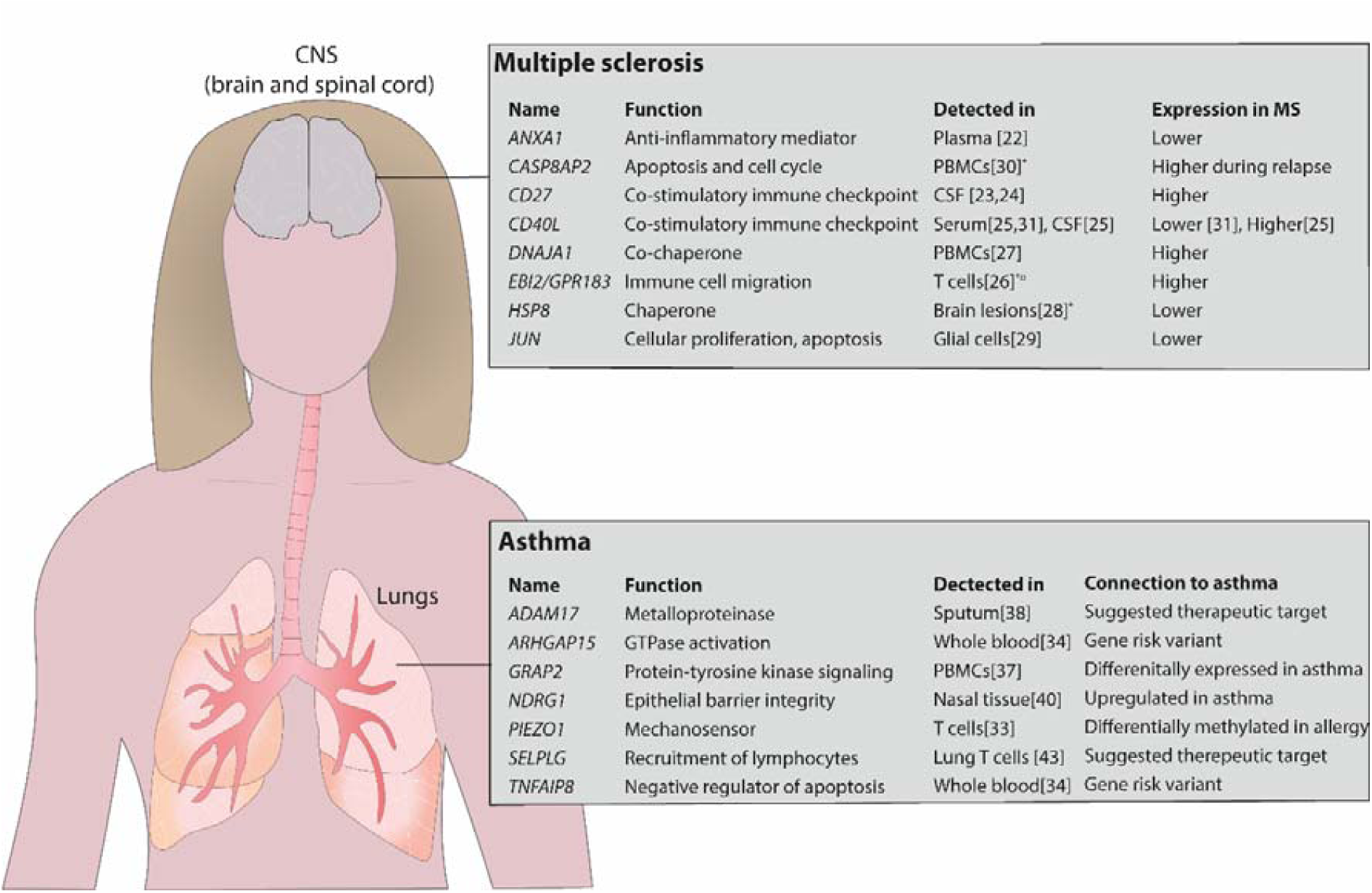
Overview of detected potential biomarkers in asthma and MS. The model identified several proteins that have previously been identified in MS and asthma. The upper panel shows the potential biomarkers identified in MS and the lower panel shows the same in asthma. *mRNA expression, ¤ identified in mice. PBMCs, peripheral blood mononuclear cells. References are given in the figure.

For asthma we found six of the top 20 genes that were differentially expressed to previously be reported for the disease (**Table S3**). Next, we analysed asthma genes uniquely identified by our model and found seven genes that had previously also been reported to be associated with disease[33-38] and are currently being evaluated as potential therapeutic targets (**Figure 4**; **Table S4).** Examples of those genes include *NDRG1*, which regulates Th2 differentiation, a key driver in asthmatic disease, downstream of the mTORC2 complex[39, 40], *ADAM17*, a metalloproteinase involved in lung inflammation[35], *PIEZO1*, a mechanosensor regulating T cell activation[41] and pulmonary inflammatory responses[42], and the P-selectin ligand encoding gene *SELPLG*, important for recruitment of lymphocytes to the airways[43, 44]. Furthermore, the immunomodulatory genes *TNFAIP8* and *ARHGAP15* were identified in GWAS studies as shared risk variants for several IgE-mediated diseases including asthma, allergic rhinitis and atopic eczema[34]. Thus, we have validated that our model can identify important biomarkers and therapeutical targets also in the context of another immune-mediated disease, *i.e.* asthma.

## Discussion

In the present study we have shown that simple mRNA-protein models, in which the protein expression is defined as a linear combination of the splice variants of a gene with a time-delay accounting for the dynamical effect induced by post-transcriptional processes and protein synthesis, can profoundly improve our ability to predict protein abundance from mRNA abundance. Furthermore, we demonstrated the impact that this finding can have within genome medicine by predicting and validating biomarkers for MS and asthma.

Despite being part of the central dogma and of uttermost importance in biology and medicine, the prediction of protein levels from mRNA levels has long been associated with low precision, which has been a matter of debate[4]. Due to the complex process of mRNA-to-protein translation, there are several aspects that need to be considered[8]. In this paper we thoroughly addressed two presumed main aspects; (1) how to incorporate splice variants into the prediction protein expression, and (2) how to deal with the time-delay of the translation between mRNA and protein expression. Interestingly, both aspects were found to impact prediction of protein abundance, as shown in our combined model, although the incorporation of splice variants influenced the protein abundance prediction the most. Herein, we report splice variants to have a wider correlation profile, both positive and negative, than what would be expected, and our novel approach takes advantage of this anti-correlation between splice variants and proteins. In previous work, the impact of incorporating splice variants into protein predictions has been analysed. These studies have focused on mechanistic cell-type independent factors such as splice variant-specific degradation rates[45]. Instead, we found that the correlations were cell-type specific and we constructed data-driven predictive models. In order to construct those models, we performed activation of NT_H_ cells followed by time-series analysis, which enabled us to infer the system based on its dynamics. These models were simplistic linear and time-delayed and validated through low out-of-sample prediction error. We found that usage of these models in complex disease enabled identification of more differentially expressed genes, which we therefore predicted as potential biomarkers. One such protein was validated as a biomarker for the MS disease prognosis. Thus, a main biological message is that intra-gene splice variant expressions influence translation, but the multifaceted nature of this mechanism remains too complex to capture with linear regression models.

Although incorporating splice variant information into the model was the main influential factor on the correlation, time delay also had an impact. The kinetics in translation of mRNA to protein is of general interest given its crucial importance in the design of experiments, for example in verifying relevance of mRNA expression to protein expression. Given that time-series experiments are time- and labor intensive, as well as expensive, a database that provides the relevant time delay between mRNA expression and the expression of its corresponding protein would be immensely valuable. Here, we present such an atlas, comprising almost 5000 gene expression-to-protein translation kinetics (**Table S1**).

A limitation with the paper is that we investigated few cell types, namely T_H_1 cells, T_REG_ cells and B cells. We also only performed wet lab experiments in one of these cell types, but were able to transfer the approach to two other cell types *in silico*, showing the robustness of the model assumptions. Furthermore, the chosen cell types are central in regulation of immune responses, and the T_H_ cells indeed are involved in many complex and common illnesses, like infectious, allergic, autoimmune and cardiovascular diseases and cancer.

In conclusion, we have constructed data-driven linear models incorporating splice variant information and time delay to with high accuracy predict protein expression from mRNA expression. We have shown the general applicability of our approach by developing models for datasets from several cell types and shown the robustness of our approach. In addition, the general principle of the model should be applicable to other cell types and can be used when that data becomes available. However, our data show that the model should be applied in a cell-specific manner given the low correlation in mixed tissue samples. We expect this modelling strategy to be generally applicable to other cellular differentiation systems, such as embryonic stem cell differentiation, and to be increasingly useful for understanding basic biology and identification of new biomarkers as more RNA-Seq and proteomic data sets become publicly available. Finally, we have shown that approach is of clinical relevance through applying it to predict validated biomarkers.

## Data availability statement

The raw and processed RNA-seq data was submitted to the EMBL-EBI sequencing archive arrayexpress and is available under the accession number <u>E-MTAB-7775</u>. The proteomics data was submitted to the EMBL-EBI proteomics repository PRIDE under the accession PXD013361.

## Ethics consent and permissions

The study was approved by the Regional Ethics Committee in Linköping, Sweden (Dnr M180-07 and M2-09). All patients were recruited at the Department of Neurology, Linköping, University Hospital Sweden and both patients and controls gave written consent prior to inclusion.

## Competing interests

The authors declare that they have no competing interests.

## Author contributions

MG initiated and supervised the study. RM and OR performed bioinformatics analyses, and RM performed the modelling. These analyses were led by MG, CA, JT, and DGC. OR performed experimental work on T-cell differentiation, which were supervised by CEN, MCJ, JE and MB. MJK and CHN performed the proteomics analysis, which was supervised by MSK. FP and JM recruited patients and collected clinical material, and SH performed and analysed the biomarker validation assays, which were led by IK, MCJ, and JE. All authors contributed to and approved the final draft for publication.

