## Supplementary material for "A validated strategy to infer protein biomarkers from RNA-Seq by combining multiple mRNA splice variants and time-delay": Suplemetary materials and methods

Supplementary information

Material and methods

**Isolation of CD4^+^ T helper (T_H_) cells and T_H_1 differentiation**

Peripheral blood mononuclear cells (PBMC) were isolated from blood donor derived buffy coats through gradient centrifugation over Lymphoprep (Axis shields diagnostics, Dundee, Scotland). Naive CD45RA^+^ CD4^+^ T cells were subsequently isolated with magnetic bead separation using a Naive CD4^+^ T Cell Isolation Kit II (Miltenyi Biotec, Bergisch Gladbach, Germany). The cells were then activated and differentiated towards T_H_1 using Dynabeads™ Human T-Activator CD3/CD28 (1 bead/cell) (Dynal AS, Lillestøm, Norway), 5 ng/µl recombinant human IL-12p70, 10 ng/µl recombinant human IL-2 and 5 µg/µl Mouse Anti-Human Il-4 Monoclonal antibody (clone MAB204) (all three from, Bio-Techne, Minneapolis, USA). All primary CD4-T-cell cultures were cultured and differentiated at 37 ֯C, with 5% CO_2_ in RPMI 1640 media containing L-glutamine, 10% FBS and 1% Penicillin/Streptomycin mixture (All from Gibco, Paisley, United Kingdom).

**Establishment of RNA-seq Time series**

To establish the earliest informative time point for the upcoming RNA-Seq, Naive CD45RA^+^ CD4^+^ T-cells were isolated and differentiated as above with sampling at 3, 5, 10, 15, 30 and 60 minutes. At each sampling point samples were washed twice with PBS by centrifugation at 1000g for 1 minute followed by snap-freezing in a dry-ice ethanol bath and transferee to -80 ֯C for storage. RNA was then extracted with a RNeasy mini kit (Hilden, Germany) and first strand cDNA synthesis performed with TaqMan™ Reverse Transcription Reagents. The cDNA was then mixed with TaqMan™ Fast Advanced Master Mix and FAM labelled TaqMan probes and primers. The qPCR assay was carried out on a 7900HT Fast Real-Time PCR System instrument (Applied biosystems) in 20 µl on a 96 well plate with probes for *IL2*, *IFNG*, *TBX21*, *GUSB* and ACTB. GUSB and ACTB were used as reference genes. All cDNA and TaqMan reagents were purchased from Applied Biosystems (Foster city, CA, USA). All probes and primers were purchased from ThermoFisher scientific (Waltham, MA, USA). qPCR was performed in technical triplicates for each time point and probe and the experiment was repeated six times (one repeat lacked the time points for 3 and 10 minutes). Expression of *IL2* and *IFNG* was significantly increased at 30 and 60 minutes (P < 0.05, students T-test) (Fig S1, *TBX21* expression not shown as it was constant). Based on this the earliest time point for the RNA-seq was set to 30 minutes.

**RNA-seq and proteomics sample collection**

Naive CD45RA^+^ CD4^+^ T-cells were isolated, differentiated and sampled as above at baseline, 0.5h, 1h, 2h, 6h and 24h for RNA-seq, and baseline, 1h, 2h, 6h, 24h and 5 days for proteomics. RNA was isolated using a ZR-Duet DNA/RNA kit (Zymo Research, Irvine, USA) and stored at -80°C until transport. During the protein extraction, multiple samples were pooled from twelve different individuals to reach the necessary amount of material for the subsequent analysis steps. The RNA and proteomics samples were not paired.

**RNA-seq sample preparation and sequencing**

RNA library preparation and the subsequent RNA-sequencing were carried out by the Beijing Genomics Institute (https://www.bgi.com/global/). Library preparation was performed using the TruSeq RNA Library Prep Kit v2 (Illumina, San Diego, USA). Each sample was sequenced to the depth of 40 million reads per sample (Fig 1A) with pair end sequencing and a read length of 100bp on an Illumina 2500 instrument.

**Proteomics sample preparation**

**Chemicals**

Urea, Ammonium bicarbonate (ABC), Triethylammonium bicarbonate (TEAB), Iodoacetamide (IAA), Ammonium formate, Trifluoroacetic acid (TFA) and formic acid were purchased from Sigma-Aldrich (St. Louis, MO). Dithiothreitol (DTT) were purchased from Roche (Hoffmann, Switzerland). Protein assay kit (# 23235) and TMT 6 plex isobaric labeling kits were purchased from Thermo Scientific (MA, USA) and MS grade Trypsin was purchased from Promega (Madison, WI).

**Cell lysis**

The isolated cells from patients were resuspended in 100 μl of 8 M Urea in 40 mM Tris-HCl (pH 7.6) and pooled with a similar number of total cells between biological replicates. The suspension was sonicated using focus sonicator (Sonic Dismembrator 500, Fisher Scientific) for 3 cycles of 10 s pulse with 10 s intervals at 10% of power. After sonication, a magnetic rack was used to remove the T-Activator beads used for the polarization. Protein concentration was measured using BCA assay and 40 μg of each sample was digested with trypsin using In-solution digestion.

**In solution digestion**

First reduction and alkylation of disulfide bonds on proteins were carried out using 1 M DTT, final sample concentration 10 mM, for 45 minutes and 1 M IAA, final sample concentration 30 mM, for 30 minutes in a dark, respectively. Following alkylation and reduction the samples were diluted with Ammonium bicarbonate buffer (pH 8.0) until the urea concentration was 1 M. The proteins were digested with trypsin overnight at 37 ℃ at an enzyme to protein ratio of 1:20. Finally, the peptides were acidified with 100 % Trifluoroacetic acid (TFA) to a final concentration of 1% TFA and then desalted using macro spin columns (Harvard apparatus, USA).

**TMT labeling**

Peptides were labeled with 6-plex TMT reagent using manufacturer’s protocol with some modification. The six peptide samples from each time series was resuspended in 100 μl of 100 mM TEAB buffer (pH 8.0) and a unit of each TMT reagent was resuspended in 40 μl of acetonitrile. Subsequently, the prepared TMT reagent was transferred to the peptide sample and then vortexed. Next, the samples were incubated for 2 hours at room temperatures. Finally, all of the labeled peptide samples from each time series were pooled and concentrated by vacuum centrifugation. The labeled sample was resuspended 100 μl with 10 mM ammonium formate in water (pH 10) and subjected to high-pH reverse-phase liquid chromatography fractionation.

**High pH Fractionation**

The TMT labeled peptides were fractionated into 24 fractions. Peptide samples were separated using an analytical column (Xbridge, C18, 5 μm, 4.6 mm x 250 mm) on the Agilent 1200 series HPLC system. Peptides were eluted using following gradient over 115 min : 0-10 min, 0% B, 10-20 min, 5% B, 20-80 min, 35% B, 80-95 min, 70% B, 95-105 min, 70% B, 105-115 min, 0% B; 10 mM ammonium formate (pH 10) was mobile phase A, and 10 mM ammonium formate with 90% ACN (pH 10) was mobile phase B. The 96 fractions were added up into 24 fractions by combining Fraction 1 - Fraction 25 - Fraction 49 - Fraction 73, Fraction 2 - Fraction 26 - Fraction 50 - Fraction 74...Fraction 24 - Fraction 48 - Fraction 72 - Fraction 96. All fractions were vacuum dried and stored at -80℃ after the desalting step.

**LC-MS analysis**

The fractionated peptides were analyzed on an Orbitrap Fusion Lumos Tribrid Mass Spectrometer (source) coupled with the Easy-nLC 1200 nano-flow liquid chromatography system (Thermo Fisher Scientific). The peptides from each fraction were reconstituted in 0.1% formic acid and loaded on an Acclaim PepMap100 Nano-Trap Column (100 μm × 2 cm, Thermo Fisher Scientific) packed with 5 μm C18 particles at a flow rate of 5 μl per minute. Peptides were resolved at 250-nl/min flow rate using a linear gradient of 10% to 35% solvent B (0.1% formic acid in 95% acetonitrile) over 95 minutes on an EASY-Spray column (50 cm x 75 µm ID), PepMap RSLC C18 and 2 µm C18 particles (Thermo Fisher Scientific), which was fitted with an EASY-Spray ion source that was operated at a voltage of 2.3 kV. Mass spectrometry analysis was carried out in a data-dependent manner with a full scan in the mass-to-charge ratio (*m/z*) range of 350 to 1,800 in the “Top Speed” setting, three seconds per cycle. MS1 and MS2 were acquired for the precursor ions and the peptide fragmentation ions, respectively. MS1 scans were measured at a resolution of 120,000 at an *m/z* of 200. MS2 scan was acquired by fragmenting precursor ions using the higher-energy collisional dissociation method and detected at a mass resolution of 30,000, at an *m/z* of 200. Automatic gain control for MS1 was set to one million ions and for MS2 was set to 0.1 million ions. A maximum ion injection time was set to 50 ms for MS1 and 100 ms for MS2. Higher-energy collisional dissociation was set to 35 for MS2. Precursor isolation window was set to 0.7 *m/z*. Dynamic exclusion was set to 35 seconds, and singly-charged ions were rejected. Internal calibration was carried out using the lock mass.

**Peptide and protein identification**

The obtained data were analysed using MaxQuant (v 1.6.0.1). MS raw data were searched using Andromeda algorithm with matching to the Uniprot human reference (released in Nov, 2017). A specificity of trypsin was determined at up to 2 missed cleavages. In modification, carbamidomethylation, TMT 6-plex modification at lysine and N-termination were set as the fixed modifications, and oxidation of methionine was set as a variable modification. The false discovery rate (FDR) for peptide level was evaluated to 0.01 for removing false positive data. For highly confident quantifications of protein, protein ratios were calculated from two or more unique quantitative peptides in each replicate. Data was normalized and removed contaminant and razor peptide. To enrich differentially expressed proteins (DEPs), we analysed the quantitative ratios (as the Log2 value). The fold-change ratio cut off was more than 2 or less than 0.5 based on intensity of 0 min. Searched data went through statistical process with Perseus (v 1.5.1.6). To interpret the expression pattern of each protein in biological pathway, biological ontology searches were performed. Biological processes and KEGG pathway were validated with DAVID bioinformatics resources^1^. Every result required a p-value less than 0.05. Enriched DEPs were used for gene ontology searches, and gene ontology information was used to draw networks among DEPs using the STRING database (http://string-db.org/).

**Mathematical modelling**

**RNA-seq data analysis**

All RNA-seq data were processed using the following pipeline. Sample qualities were assessed with fastQC and the mRNA reads were subsequently aligned using STAR^2^ to the “Homo_sapiens.GRCh37.75.dna.primary_assembly.fa” from Ensemble. The sample reads contained no sequencing adapters and sequence quality was very high so no cutting of adapters or filtering of low quality reads were performed prior to alignment. The resulting read alignment bam files were assembled into transcripts with StringTie^3^ using the GRCh37.75 gtf annotation from Ensemble. To evaluate mRNA to protein interactions, mRNA reads were mapped to the mass spectrometry signal of protein abundance using the Homo.sapiens and Mus.musculus package in R^4^. Next, correlations were determined using Pearson correlations, as calculated using the SciPy-library for python 3.6^5,6^. All gene-protein correlations that could be mapped were used to calculate the mean correlation coefficient. This was done by considering each gene and protein time series individually, *i.e.* calculating one correlation per gene.

**Model construction**

We hypothesized that protein abundance could be predicted using a linear combination of the corresponding splice variants. To predict protein abundance, we used the Sklearn^7^ implementation of the LASSO^8^, an L1-penalized linear regression model.

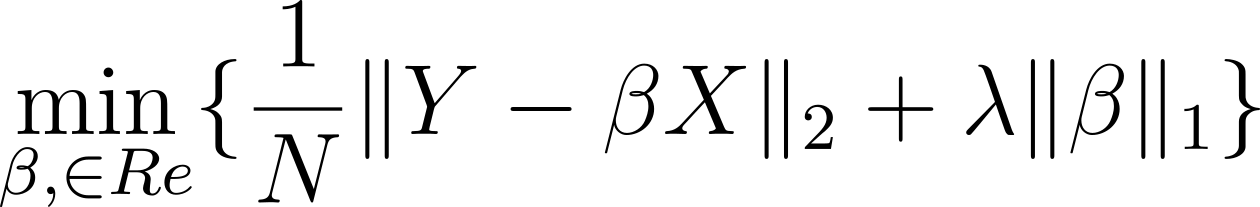

Here, the time series of one protein is denoted the vector Y, and the corresponding time series of the splice variants are denoted by the matrix X. The rate constant for each splice variant is contained in the vector β. Furthermore, the λ parameter regulates the influence of the L1 term, and was determined individually for each protein. The λ term was chosen to minimize the prediction error of a leave-one-out cross validation. In the Th1 dataset, the time points differed such that the mRNA abundance also had a measurement at t=30 minutes, while the protein data instead had a measurement of t=120h. For comparison, the protein data for 30 minutes was interpolated, while the 120h time point was omitted. The same procedure was performed using the regulatory T cells from Schmidt *at al*^9^ were Treg induced by either TGF-β, TGF-β and ATRA, or TGF-β and butyrate. Lastly, the same procedure was performed for mice B-cells were B-cell differentiation was induced by the Ikaros transcription factor (GSE75417). Pipe-line and code available from

<https://gitlab.com/Gustafsson-lab/IMUNA-an-integrated-multilevel-Th1-analysis> .

**Time delay analysis**

The effect of time delays between mRNA and protein was analyzed since this might affect the prediction of protein abundance. First, we considered the Th1 data and linearly interpolated 400 data points between 0 and 24h for both the mRNA expression and protein abundance data. Next, we calculated the correlation between each pair of interpolated time points, i.e. in total 160 000 comparisons. From this analysis we observed a mean? time delay of approximately 6 hours. These results prompted us to include a time delay in our model of splice variants to predict protein abundance.

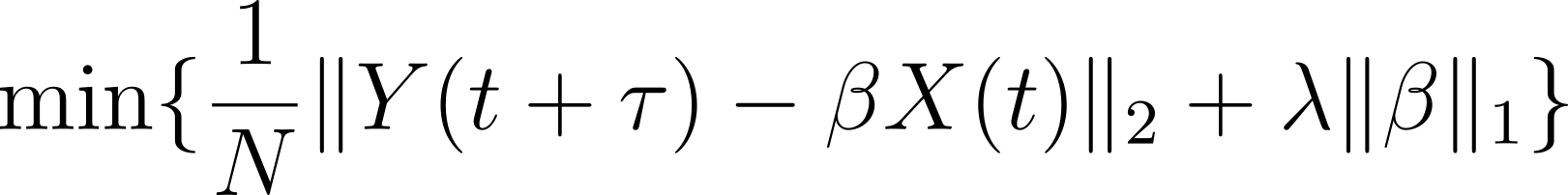

In the updated model, we added a protein specific time delay τ to regulate which time point of splice variant expression should be used. As an example, a τ = 0.5h would result in splice variant abundance of t = [0h,1h,2h,6h,24h] predict protein abundance interpolated at t = [0.5h, 1.5h, 2.5h, 6.5h, 24.5h]. For each τ, we did a leave-one-out cross validation on top of the one used for determining the λ parameter. Next, the τ yielding the lowest cross validation error was selected.

**Disease prediction**

Disease relevance of the splice variant models was tested by re-analysis of deep RNA-sequenced case control material of samples containing total CD4 + T-cells, i.e. CD4 + T-cells with all its sub-types. We found T-cell prolymphocytic leukemia (T-PLL, GSE100882), asthma in obese children (GSE86430), and allergic rhinitis/asthma (GSE75011) studies through a Gene Expression Omnibus (GEO) repository search and multiple sclerosis (MS) through collaboration^10^. For each of the studies’ datasets we used the T_H_1 and Treg derived models on how to combine mRNA splice variants to predict protein abundance. The resulting sets of predicted protein levels were tested for differential expression between patients and controls using a non-parametric Kruskal-Wallis test. We also applied Kruskal-Wallis tests to the individual splice variants that were used by the models. We assessed model effects by measuring the increase of nominally differential expression from model predictions compared to ingoing splice variants into the model.

**Protein validation**

**Patients and controls**

Cerebrospinal fluid (CSF) was collected from a cohort of 41 patients with newly diagnosed clinically isolated syndrome (CIS) or relapsing remitting MS (RRMS) (table S5) that has been described in more detail elsewhere^11^. All patients fulfilled the revised McDonald criteria from 2010^12^. The patients were followed, and new samples obtained after one, two and four years. Disease activity was assessed using “no evidence of disease activity” (NEDA), defined by no clinical relapses, no sustained EDSS progression and no new T2 or Gadolinium enhancing lesions. 12 patients at the two year- and 7 patients at the four-year follow-up were classified as NEDA, whereas patients with relapses, brain MRI activity and sustained disease progression were classified as “evidence of disease activity” (EDA; n=27 and n=32 at two and four years, respectively). Two patients did not complete the study^11^. Twenty-three healthy age-and sex-matched blood donors were included as controls. A second cohort of CSF samples from 16 Natalizumab-treated patients with RRMS or secondary progressive MS (SPMS) was also included. CSF samples were obtained (out of a total of ≈70 included patients with RRMS or SPMS) before and after one year of treatment with Natalizumab were also included in the study (table S5). This study cohort has been described previously^13–15^. All patients were recruited at the Department of Neurology, Linköping, University Hospital Sweden and both patients and controls gave written consent prior to inclusion. The study was approved by The Regional Ethics Committee in Linköping.

**Protein measurements**

Quantification of sCD27 was performed using the Human Instant ELISA™ kit from eBioscience (Thermo Fischer Scientific, Waltham, MA, USA) according to the instructions provided by the manufacturer. The optical densities (O.D.) were read at 450 nm with a wavelength correction at 620 nm in a Sunrise™ microplate reader (Tecan, Shanghai, China). Data acquisition was performed using Magellan™ version 7.1 computer software (Tecan). The lowest detection limit was 0.63 U/ml and values below the detection limit were given half the value of the detection limit. Statistical differences were determined using Mann-Whitney U-test or Wilcoxon matched-pairs signed rank test (Graphpad Prism v7.04, San Diego, CA, USA) Annexin A1, measured by the human Annexin A1 ELISA kit (Abcam, Cambridge, United Kingdom), was undetectable in all analyzed samples (n=32, of whom n=16 were included before and 16 after one year of treatment with Natalizumab). Multiplex Bead Technology (MILLIPLEX® MAP Kit, Cat. #: HCYTOMAG-60K-01, Merck Millipore, Burlington, MA, USA) was used to measure soluble CD40L according to the manufacturer's description. The samples were analysed on a Luminex®200™ instrument (Invitrogen, Carlsbad, CA, USA) and data was collected using xPONENT 3.1™ (Luminex Corporation, Austin, TX, USA) analysed using the MasterPlex® Reader Fit (MiraiBio Group, Hitachi Solutions America Ltd, San Bruno, CA, USA). The lowest detection limit was 1.6 pg/ml and values below the detection limit were given half the value of the detection limit. sCD40L concentration was below the lowest detection limit in 71 out of 96 samples (74% undetectable) and was therefore considered as undetectable.

Supplementary Figures

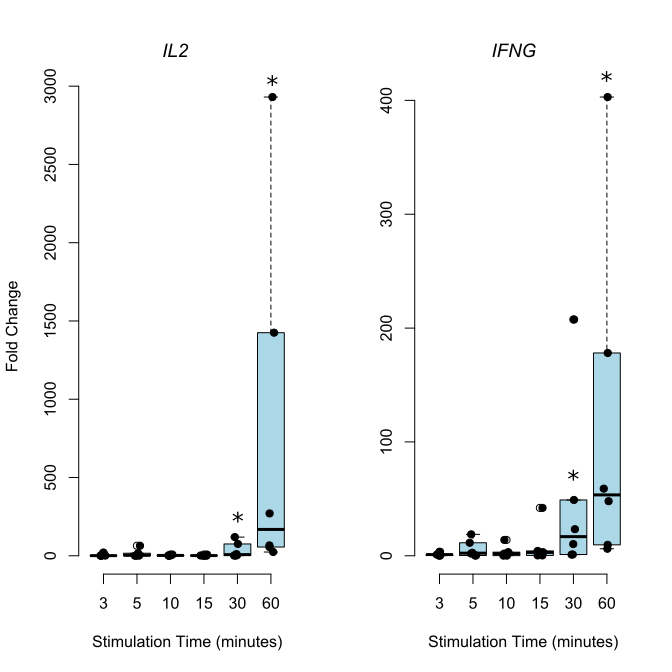

**Figure S1: In vitro T_H_1 differentiation from naïve T cells resulted in significant consistent upregulation of** *IL2* and *IFNG* expression observed at 30 minutes and at 60 minutes (p < 0.05, student’s t-test) as measured by qRT-PCR.

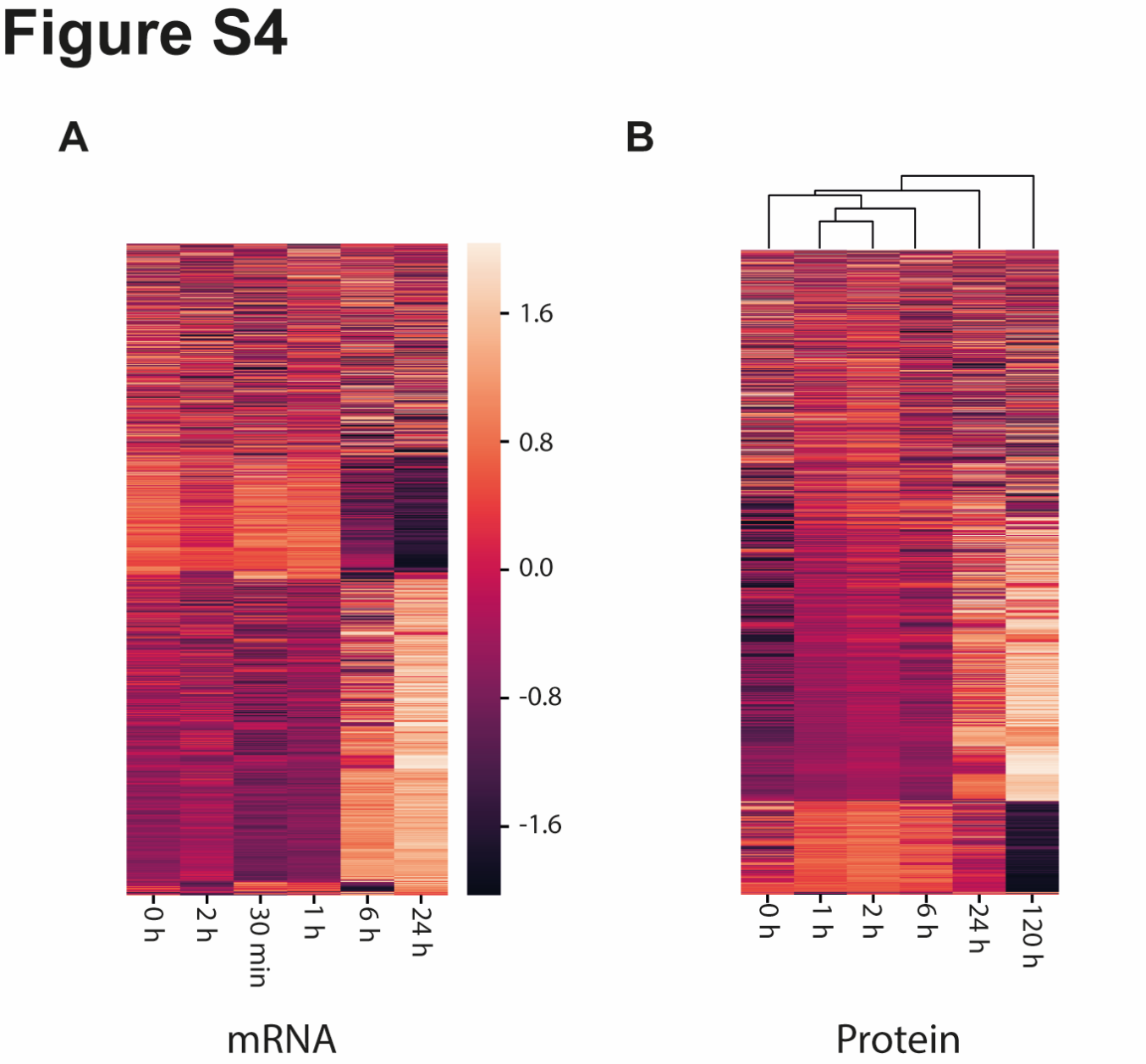

**Figure S2. Hierarchichal clustering of time-series RNA-seq (A) and mass-spectronomy proteomics data of Th1 differentiation showed that samples clustered well with respect to time.** We analysed the data quality, and the similarities of the different measured time points, by hierarchical clustering. For all 4860 genes and corresponding protein time series that were used for the model estimation, we performed a Z-normalization of each time series and performed a hierarchical clustering using the Python package Scipy. We used the default parameters ‘single linkage’, minimizing the Euclidian distance between clusters. In A), we clustered the mRNA gene expressions and found all biological repeats except the time point t=2h to cluster with their closest neighbours in time. In B), we saw a perfect clustering of the time point in a consecutive order. The clustering in A) and B) show that there is a distinguishable biological signal of activation throughout both the mRNA expression and protein abundance datasets.

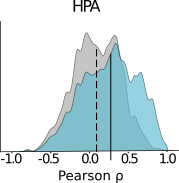

**Figure S3. Gene/protein correlations in data from the human protein atlas.** In the histogram, the grey curve shows the correlation distribution when the sum of all splice variant expressions of a transcript is used to quantify mRNA abundance (median: dashed line at 0.21), while in the blue histogram our multiple splice variant based model is used. Only cross-validated protein predictions are shown for the 3410 out of 4920 proteins for which the null-model could be rejected.

Supplementary files

**Table S1: Table of all genes with their transcripts and modelling coefficients across all three models.** Each sheet represents one cell-type, namely human T_H_1, T_REG_, and mice B-cells, were each row represent one transcript to protein association (see section time-delayed models for details). Column A is the gene name of the corresponding protein, column B the splice variant name, column C the value of the linear coefficient, column D is the correlation between measured and predicted protein abundance values, column E is the corresponding time-delay (τ).

**Table S2: Top 20 differentially expressed proteins in multiple sclerosis.** Markers previously associated with multiple sclerosis are indicated in grey.

**Table S3: Top 20 differentially expressed proteins in asthma.** Markers previously associated with asthma are indicated in grey.

**Table S4: Predicted differentially expressed proteins not detectable by standard RNA-Seq data analysis.** Markers previously associated with asthma are indicated in grey.

**Table S5: Patient and healthy control characteristics at inclusion and follow-up**
